# Grid partitioning image analysis for bacterial cell aggregates

**DOI:** 10.1101/2025.05.22.655429

**Authors:** Yuki Ohara, Shogo Yoshimoto, Katsutoshi Hori

## Abstract

Bacterial cell aggregation plays a fundamental role in surface colonization, stress tolerance, and interspecies metabolite exchange; yet quantitative analysis remains challenging. Here, we introduce grid partitioning image analysis (GPIA), a simple workflow that quantifies the compositional heterogeneity of bacterial aggregates. Confocal laser scanning microscopy (CLSM) images of fluorescently labeled *Acinetobacter* sp. Tol 5, which exhibits a self-aggregative nature through its cell surface protein AtaA, were partitioned into 2-µm square grids. Grids containing one or no cells were classified as dispersed, whereas those containing multiple cells were classified as aggregates, and the proportion of EGFP-labeled cells within each grid was recorded. Reference images representing dispersed cells, homo-aggregates, and hetero-aggregates produced characteristic EGFP-ratio histograms that matched binomial predictions. When AtaA production in one cell type was decreased, the histogram changed from a symmetric unimodal histogram with the peak at 40–60 % EGFP-ratio to a skewed distribution, indicating that GPIA can detect differences in cell-to-cell affinity. Using the same procedure, we examined six in-frame deletion variants of AtaA. The deletion of the N-terminal head domain alone prevented co-aggregation with full-length AtaA, suggesting that homophilic recognition by this domain mediated self-aggregation, whereas deletions in all other regions had no measurable effect. GPIA, therefore, offers a simple and rapid approach for quantitative studies on bacterial cell aggregation, bridging the gap between qualitative microscopy and quantitative but technically demanding single-cell analysis. GPIA will accelerate research on cell–cell interactions, which are the foundational processes that drive biofilm formation and the assembly of microbial consortia.

## 1. Introduction

Bacterial cell aggregation, which includes homotypic aggregation of identical cells and heterotypic aggregation among different types of cells, plays a fundamental role in microbial ecology and pathogenesis (Nwoko and Okeke, 2021; Kragh et al., 2023). By facilitating close cell-to-cell contacts, aggregation promotes initial surface colonization, shields communities from shear stress, desiccation, and antimicrobials, and accelerates interspecies signaling and metabolite exchange (Kruse et al., 2021; Liu et al., 2024). These cell-cell interactions are mediated by cell-surface adhesins, such as extracellular polysaccharides and protein fibers (Formosa-Dague et al., 2016; Trunk et al., 2018). In the medical field, bacterial cell aggregation is treated as a nuisance (Nwoko and Okeke, 2021; Liu et al., 2024). On the other hand, it is recognized as beneficial in wastewater treatment and bioprocesses for chemical production to concentrate biomass and stabilize the systems (Sethi et al., 2023; Najim et al., 2024; Hammond et al., 2025). Therefore, understanding the characteristics and mechanisms of bacterial cell aggregation is important in various research fields.

*Acinetobacter* sp. Tol 5 shows remarkable nonspecific adhesiveness to various material surfaces and a self-aggregative nature through its cell surface nanofiber protein AtaA (Ishikawa et al., 2012). AtaA is a member of trimeric autotransporter adhesins (TAAs), which are outer membrane (OM) proteins widely distributed in gram-negative bacteria (Leo et al., 2012). The polypeptide chains form a homo-trimeric structure with an N-terminal passenger domain (PSD) corresponding to its adhesive functions and a C-terminal transmembrane domain that transports and anchors the PSD onto the OM (Bassler et al., 2015). The adhesive and aggregation features of AtaA can be conferred to other non-adhesive gram-negative bacteria by transformation with the *ataA* gene (Ishikawa et al., 2014; Yoshimoto et al., 2023). Previously, we invented a new method for immobilizing bacterial cells utilizing AtaA and demonstrated its advantages: large numbers of bacterial cells expressing AtaA can be quickly immobilized onto various material supports, the immobilized cells can be reversibly detached by rinsing with deionized water or by adding casein hydrolysates, and the immobilized cells can be efficiently used for bioproduction (Yoshimoto et al., 2017; Ohara et al., 2019; Usami et al., 2020). Although AtaA-mediated self-aggregation plays an important role in the initial attachment of bacterial cells to the material surfaces and in increasing the number of immobilized cells by stacking and flocculation (Furuichi et al., 2020), the details of aggregation have not been investigated.

In microbiology, self- and co-aggregation are commonly assessed by simple tube-settling assays that monitor turbidity (Trunk et al., 2018; Nwoko and Okeke, 2021), but the measurement of co-aggregation between self-aggregative bacteria is difficult. Microscopy-based evaluations of bacterial aggregation have been reported, but they remain largely qualitative, and the few quantitative image analysis methods reported to date are technically complicated (Glass and Riedel-Kruse, 2018; Khalil et al., 2020). In contrast, eukaryotic cell–cell adhesion has been evaluated by simpler image analysis, yielded quantitative descriptions of the interaction (Sieber and Roseman, 1981). Transferring such simple quantitative imaging strategies to bacteria is beneficial for rapid and intuitive analysis; however, it requires extensive modifications and optimization because bacterial cells are much smaller than eukaryotic cells and form densely packed clusters that obscure individual cell boundaries and confound automated segmentation.

In this study, we developed a new method of grid partitioning image analysis (GPIA) that quantifies the compositional heterogeneity of bacterial aggregates (Fig. 1). Using Tol 5 derivatives that display AtaA and distinct fluorescent reporters, we demonstrate that GPIA (i) distinguishes between aggregated and dispersed cells, (ii) evaluates the heterogeneity of cell aggregates, and (iii) detects the changes in interaction affinity caused by modulating the production level of AtaA and the in-frame deletion of AtaA.

**Figure 1.**
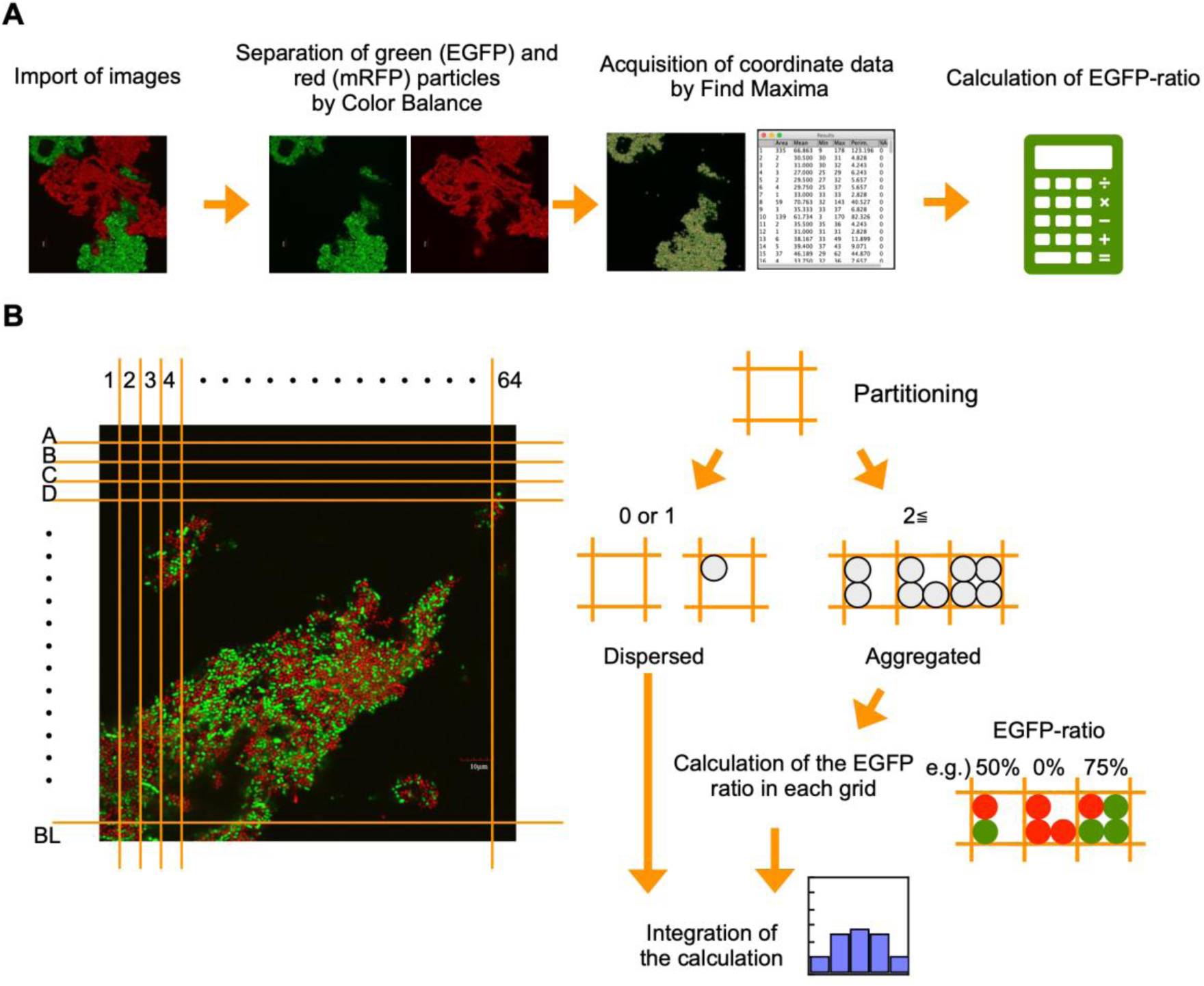
Overview of grid partitioning image analysis (GPIA). (A) CLSM images were imported into ImageJ and separated into EGFP and mRFP channels using the “Color Balance” function. Fluorescent cell coordinates were extracted independently from each channel using the “Find Maxima” function, and the resulting coordinate sets were exported as a spreadsheet for further analysis. (B) Concept of grid partitioning and cell classification. Based on the imported cell coordinates, the image was divided into 64 × 64 grids (2 µm × 2 µm). A grid containing one or no cells was classified as “dispersed,” while grids containing multiple cells were defined as “aggregated.” For each aggregated grid, the proportion of EGFP-fluorescent cells was calculated and binned into one of five EGFP-ratio ranges: 0 ≤ x < 20 %, 20 ≤ x < 40 %, 40 ≤ x < 60 %, 60 ≤ x < 80 %, and 80 ≤ x ≤ 100 %. The number of EGFP and mRFP cells within each bin was summed across all aggregated grids. The total number of cells across both dispersed and aggregated grids was used as the normalization baseline (100 %), and the binned values were plotted as percentage histograms.

## 2. Materials and Methods

### 2.1 Bacterial strains and culture conditions

*Escherichia coli* XL10-Gold and DH5α were used for plasmid construction. *E. coli* S-17 strain was used for plasmid conjugation (Simon et al., 1983). *E. coli* cells were grown in Luria-Bertani (LB) medium at 37°C with shaking at 150 rpm. *Acinetobacter* sp. Tol 5 (Hori et al., 2001) and its Δ*ataA* mutant strain 4140 (Tol 5 Δ*ataA*) (Ishikawa and Hori, 2013) were grown in LB medium at 28°C with shaking at 115 rpm. Antibiotics were added at the following concentrations as needed: ampicillin (100 µg/mL), kanamycin (50 µg/mL), gentamicin (10 µg/mL). L-arabinose was added to the culture medium to induce the expression of *ataA* gene on pAXG or pAXR plasmids. The concentration of arabinose varied from 0.01 % to 0.5 % to control the production levels of AtaA.

### 2.2 DNA manipulation

Plasmids used in this study are listed in Table 1. The DNA fragment encoding the enhanced green fluorescent protein (*egfp*) gene was amplified by PCR from pEGFP-C3 (GenBank: U57607.1) using primers EGFPtoC007-Fw/EGFPtoC007-Rv (Fw: CAATTAAGCTTATGGTGAGCAAGGGCGAGG, Rv: GTATTCCATGGTTACTTGTACAGCTCGTCCATGC). The plasmid backbone including the P_LtetO1_ promoter for constitutive gene expression was amplified by PCR from pC007 (Abudayyeh et al., 2016) using primers C007inv-Fw/C007inv-Rv (Fw: GTATTCCATGGTAAGGATCTCCAGGCATCAAATAAAAC, Rv: CATTAAAGCTTTTTCTCCTCTTTCAGATCCGTGC). These DNA fragments were digested with HindIII and NcoI and ligated, generating pC007G. To construct pAXG and pAXR, which encode *egfp* or *mrfp* under the P_LtetO1_ promoter (Lutz and Bujard, 1997), the DNA fragments encoding *egfp* and *mrfp* were amplified by PCR from pC007G or pC007 using primers RFPtoARP-Fw/RFPtoARP-Rv (Fw: CAATTAAGCTTATGGTGAGCAAGGGCGAGG, Rv: GTATTCCATGG TTACTTGTACAGCTCGTCCATGC). The amplified DNA fragments were assembled with PvuII-digested pARP3 by NEBuilder HiFi DNA Assembly master mix (New England BioLabs, Ipswich, MA, USA). To construct pAXG-*ataA* and pAXR-*ataA*, which were used for co-expression of *ataA* and each fluorescent protein gene, the DNA fragments encoding *ataA* genes excised from pAtaA by digestion with EcoRI and XbaI were ligated with pAXG or pAXR digested with the same restriction enzymes. The construction of the co-expression plasmids for the in-frame deletion mutant of *ataA* and fluorescent protein was conducted in the same manner using pARP3 plasmid harboring a gene encoding each in-frame deletion mutant of *ataA* (Yoshimoto et al., 2023). Transformation of the Tol 5Δ*ataA* with these expression plasmids was carried out by conjugal transfer from *E. coli* S17-1 strain (Simon et al., 1983), as previously described (Ishikawa et al., 2012).

**Table 1.** Plasmids used in this study.

| Plasmids | Description | Reference |
| --- | --- | --- |
| pC007 | Vector encoding *mrfp* under P<sub>LtetO1</sub> promoter, Amp<sup>R</sup> | (Abudayyeh et al., 2016) |
| pC007G | Vector encoding *egfp* under P<sub>LtetO1</sub> promoter, Amp<sup>R</sup> | This study |
| pEGFP-C3 | Vector encoding *egfp*, Kan<sup>R</sup> | Purchased from Takara Bio (Shiga, Japan) |
| pARP3 | Expression vector for *E. coli* and *Acinetobacter*, Gm<sup>R</sup>, Amp<sup>R</sup> | (Ishikawa et al., 2012) |
| pAtaA | Expression vector encoding *ataA* under AraC/P<sub>BAD</sub> promoter, Amp<sup>R</sup>, Gm<sup>R</sup> | (Ishikawa et al., 2012) |
| pAXG | Expression vector encoding *egfp* under P<sub>LtetO1</sub> promoter, Amp<sup>R</sup>, Gm<sup>R</sup> | This study |
| pAXR | Expression vector encoding *mrfp* under P<sub>LtetO1</sub> promoter, Amp<sup>R</sup>, Gm<sup>R</sup> | This study |
| pAXG-AtaA | Expression vector encoding *egfp* under P<sub>LtetO1</sub> promoter and *ataA* under AraC/P<sub>BAD</sub> promoter, Amp<sup>R</sup>, Gm<sup>R</sup> | This study |
| pAXR-AtaA | Expression vector encoding *mrfp* under P<sub>LtetO1</sub> promoter and *ataA* under AraC/P<sub>BAD</sub> promoter, Amp<sup>R</sup>, Gm<sup>R</sup> | This study |
| pAXG-ΔNhead | pAXG vector encoding *ataA* fragment carrying an in-frame deletion of 60-313 aa | This study |
| pAXG-ΔNS-A1 | pAXG vector encoding *ataA* fragment carrying an in-frame deletion of 327-506 aa | This study |
| pAXG-ΔNS-A2 | pAXG vector encoding *ataA* fragment carrying an in-frame deletion of 507-1337 aa | This study |
| pAXG-ΔNS-B | pAXG vector encoding *ataA* fragment carrying an in-frame deletion of 1338-2335 aa | This study |
| pAXG-ΔNS-C-ΔChead | pAXG vector encoding *ataA* fragment carrying an in-frame deletion of 2397-3169 aa | This study |
| pAXG-ΔCstalk | pAXG vector encoding *ataA* fragment carrying an in-frame deletion of 3170-3475 aa | This study |

### 2.3 Detection of co-expression of AtaA and fluorescent protein

Protein production was examined by SDS-PAGE followed by Coomassie Brilliant Blue (CBB) staining and immunoblotting using anti-AtaA_59-325_ antiserum, as described previously (Ishikawa et al., 2012).

The presentation of AtaA on the cell surface and the production of fluorescent proteins were confirmed by immunofluorescence microscopy using anti-AtaA_59-325_ antiserum, as described previously with a slight modification (Yoshimoto et al., 2023). Alexa Fluor 647 conjugate of anti-rabbit IgG (H+L), F(ab’)_2_ Fragment (Cell Signaling Technology, Danvers, MA, USA) was used for the detection of the primary antiserum. These prepared samples were observed by a confocal laser scanning microscope (CLSM; FV1000D IX81-FD/NIH, Olympus Corporation, Tokyo, Japan) with 473, 559, and 635 nm lasers.

Tube-settling aggregation assays of bacterial cells were performed as described previously (Ishikawa et al., 2012). In brief, glass test tubes containing cell suspension with an optical density at 660 nm (OD_660_) of 0.5 were left to stand at 28°C. The aggregation ratio was calculated from the decrease in the OD_660_ of the cell suspension using the following equation:

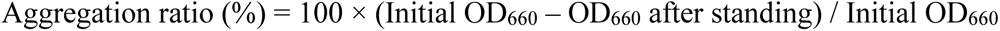

### 2.4 Formation of cell aggregation

Grown cells were transferred to a 15-mL protein low adsorption tube (Proteosave SS; Sumitomo Bakelite, Tokyo, Japan) and diluted to OD_660_ =0.5 with fresh medium. 0.5 ml of two cell suspensions containing different types of cells were mixed in a 1.5-mL protein low adsorption tube (Sumitomo Bakelite, Tokyo, Japan). For samples mixed at a 1:3 ratio, 0.25 ml and 0.75 ml of cell suspension were mixed. The mixed cell suspension was centrifuged at 5,000 ×g for 5 min, and the supernatant was discarded. The cell pellet was rinsed with deionized water and centrifuged at 5,000 ×g for 5 min. The cell pellet was re-suspended in equal volume of BS-N buffer (34.5 mM Na_2_HPO_4_, 14.7 mM KH_2_PO_4_, 15.5 mM K_2_SO_4_; pH 7.2). The cell suspension was slowly stirred at 8 rpm using a rotator (NRC-20D, Nissinrika, Tokyo, Japan) for 15 min to form cell aggregates. Prepared cell aggregates were placed onto a glass slide and observed by CLSM. The lasers of 473 and 559 nm were used for exciting the fluorescent proteins EGFP and mRFP, respectively. Confocal images were acquired using a 100×/1.4 NA oil immersion objective lens and saved at a resolution of 1024 × 1024 pixels.

### 2.5 Grid partitioning image analysis

The CLSM images were imported into ImageJ (Schneider et al., 2012). EGFP and mRFP fluorescent signals were separated using the “Color Balance” function; individual fluorescent particles were detected with “Find Maxima,” and their coordinates were exported as a spreadsheet, where the field of view was subdivided into 2 µm-square grids. The pixel scale of the CLSM image was calibrated to micrometer units. Grids containing one or no cells were classified as dispersed, while those containing two or more cells were classified as aggregated. For every grid classified as aggregated, the proportion of EGFP-fluorescent cells (EGFP-ratio) was calculated by dividing the number of EGFP-fluorescent cells by the total number of fluorescent cells (EGFP + mRFP) within that region. The grids classified as aggregated were then categorized into five bins based on EGFP-ratio: 0 ≤ x < 20 %, 20 ≤ x < 40 %, 40 ≤ x < 60 %, 60 ≤ x < 80 %, and 80 ≤ x ≤ 100 %. For each bin, the total number of EGFP and mRFP cells was summed across all grids classified as aggregated. These values were then normalized by the total number of fluorescent cells present in both grids classified as dispersed and aggregated, and the normalized percentages were plotted as histograms. The template spreadsheet used for these calculations is available in Supporting Information File S1.

### 2.6 Control conditions for theoretical distributions

To interpret experimental histograms, theoretical EGFP-ratio distributions were generated for each of the control conditions under defined assumptions. For the dispersed control, every cell was assumed to occupy its own grid, yielding no grids assigned as aggregated and therefore a flat distribution of 0 % across all EGFP-ratio bins. In the homo-aggregation control, we posited that each grid contained only a single cell type, while the overall field maintained a 1:1 mixture of EGFP- and mRFP-expressing cells; this produced a theoretical profile composed solely of the 0–20 % and 80–100 % bins, each contributing 50 % of the total. For the hetero-aggregation control, the expected distributions were modeled with the binomial formula:

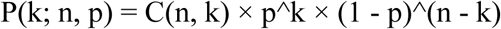

where n is the number of cells in a grid, k is the number of EGFP-fluorescent cells, and p is the probability of selecting an EGFP-fluorescent cell. We set p = 0.5 for the equal-volume (1:1) mixture and p = 0.25 for the 1:3 mixture, calculated the EGFP-ratio (k/n × 100) for k = 0–n, and assigned each outcome to the standard five bins (0 ≤ x < 20 %, 20 ≤ x < 40 %, 40 ≤ x < 60 %, 60 ≤ x < 80 %, and 80 ≤ x ≤ 100 %). This computation was repeated for n = 2–8, spanning the observed range of cells per grid. The resulting bin probabilities were combined with the weights based on the observed frequency of each grid size (Fig. 3B), generating the theoretical distributions.

## 3. Results

### 3.1 Construction of bacterial cells co-expressing *ataA* and two types of fluorescent protein genes

First, we constructed *ataA* and fluorescent gene co-expressing Tol 5 cells to distinguish the two types of cells in the image analysis. The co-expression plasmids were designed and constructed as shown in Fig. 2A and Table 1. Either *egfp* and *mrfp* genes were placed under the constitutive P_LtetO1_ promoter, while the *ataA* gene was placed under the inducible AraC/P_BAD_ promoter.

**Figure 2.**
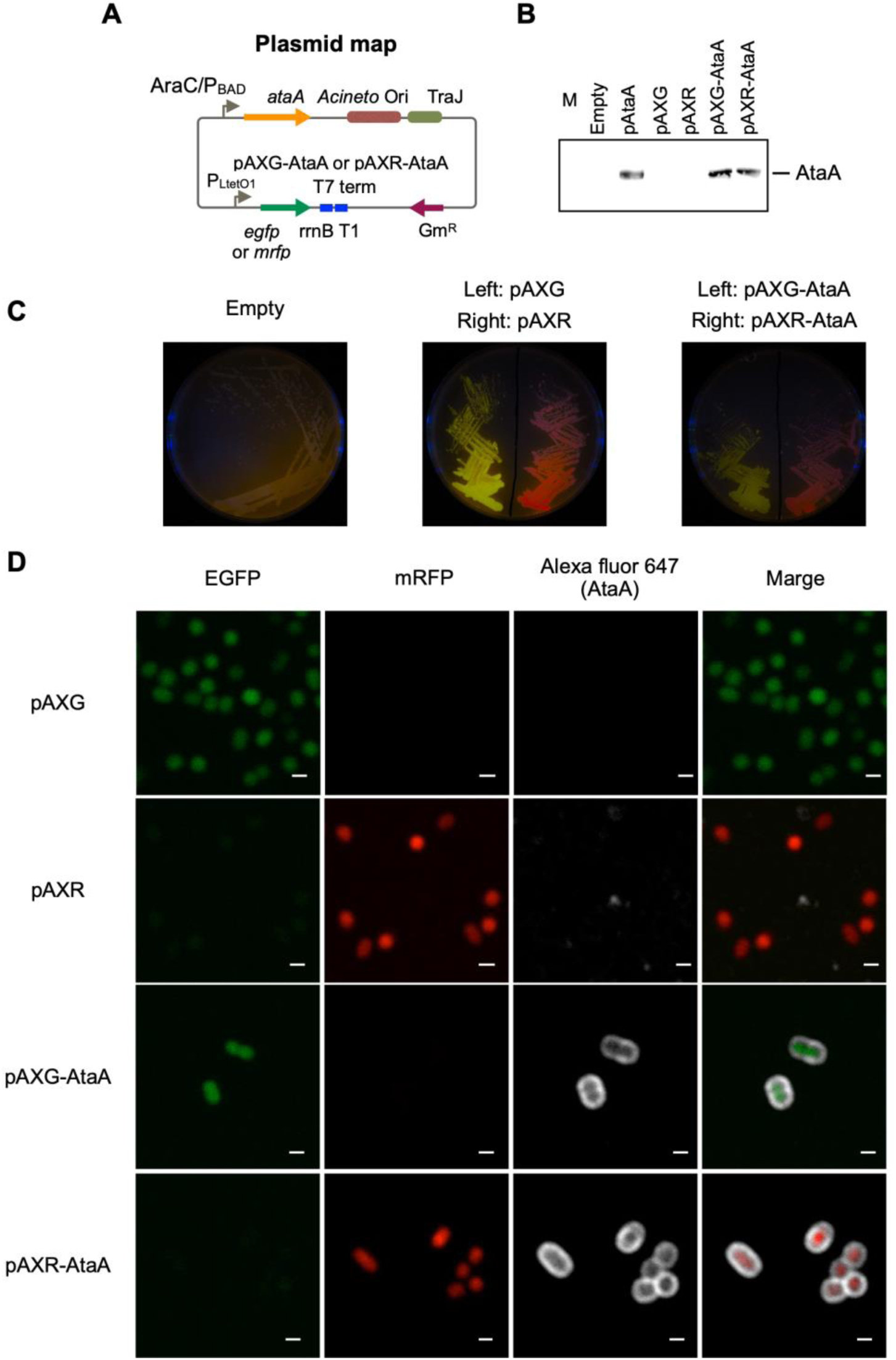
Construction of AtaA/EGFP or AtaA/mRFP co-expression strain. (A) Map of the co-expression plasmid for AtaA and fluorescent protein genes. The *ataA* gene was inserted under the AraC-P_BAD_ inducible promoter, and *egfp* or *mrfp* were inserted under the P_LtetO1_ constitutive promoter. (B) Confirmation of *ataA* expression. The whole-cell lysates of Tol 5 Δ*ataA* or its derivative mutants were analyzed by immunoblotting using anti-AtaA antiserum. (C) Photographs of agar plates taken under black light. Green and red are fluorescence derived from EGFP and mRFP, respectively. (D) CLSM observation of immuno-stained cells co-expressing *ataA* and fluorescent protein genes. Green and red are derived from EGFP and mRFP, respectively. White was derived from immuno-stained AtaA. Scale bars: 1 µm.

The constructed plasmids were introduced into Tol 5 Δ*ataA*, and the co-expression of each fluorescent protein gene and *ataA* was examined by immunoblotting and immunofluorescence microscopy. The production amount of AtaA and cell adhesiveness of Tol 5 Δ*ataA* (pAXG-AtaA) (EGFP-AtaA(+)) and Tol 5 Δ*ataA* (pAXR-AtaA) (mRFP-AtaA(+)) were almost the same as Tol 5 Δ*ataA* (pAtaA) (Fig. 2B). Tol 5 Δ*ataA* (pAXG) (EGFP-AtaA(-)) and Tol 5 Δ*ataA* (pAXR) (mRFP-AtaA(-)) exhibited intracellular fluorescence corresponding to each fluorescent protein (Fig. 2C, D). Tol 5 Δ*ataA* (pAXG-AtaA) and Tol 5 Δ*ataA* (pAXR-AtaA) simultaneously exhibited fluorescence surrounding the cells corresponding to AtaA in addition to intracellular fluorescence corresponding to each fluorescent protein (Fig. 2D). These results demonstrated that each fluorescent protein gene and *ataA* were co-expressed in Tol 5 Δ*ataA*.

### 3.2 Evaluation of cell aggregation and heterogeneity by grid partitioning image analysis (GPIA)

For the previous analysis of eukaryotic cells by Sieber and Roseman, each four-cell aggregate was treated as a single analytical unit (Sieber and Roseman, 1981). Here, we investigated the optimal subdivision size for CLSM images to divide aggregates of bacterial cells, whose size is approximately 1 µm, into units of about four cells. We divided the CLSM image of well-mixed hetero-aggregated cells shown in Figure 1B into square grids of 1, 2, and 4 µm and counted the number of cells in each grid. In 1 µm square grids, over 70 % of the cells were present individually within a single grid (Fig. S1, green). Because we defined “aggregated” as a grid containing two or more cells, 1 µm partitioning led to misclassification. In 4 µm square grids, many grids contained ten or more cells (Fig. S1, blue), so cells that were not in direct contact could be falsely classified as co-aggregated. In 2 µm square grids, most of the grids contained two to four cells (Fig. S1, orange), which were correctly classified as aggregated. These results indicate that the 2 µm partitioning is optimal for this bacterial cell. Therefore, all subsequent analyses used a 2 µm grid.

To test the performance of GPIA, four typical images were prepared by mixing the two types of AtaA-displayed (or not displayed) fluorescent cells. The cell mixture of EGFP-AtaA(−) and mRFP-AtaA(−) was prepared as the dispersed control (Fig. 3A left). The second control consisted of two cell types that can self-aggregate individually but do not interact with each other; we referred to this as the homo-aggregation control. This sample was prepared by forming homotypic cell aggregates of EGFP-AtaA(+) and mRFP-AtaA(+) cells and then mixing them (Fig. 3A middle left). The third and fourth controls consisted of two cell types that interacted with one another and with themselves to a similar extent; we designated this as the hetero-aggregate control (Fig. 3A middle right and right). These samples were prepared by mixing equal or 1:3 volumes of EGFP-AtaA(+) and mRFP-AtaA(+) cell suspensions, allowing simultaneous aggregation. These samples were observed by CLSM and analyzed according to the GPIA concept presented in Figure 1. In the dispersed control, over 70 % of cells were classified as dispersed particles (Fig. 3B, left). On the other hand, over 75 % of cells were classified as aggregates in the homo- and hetero-aggregation controls (Fig. 3B, middle left, middle right, and right).

**Figure 3.**
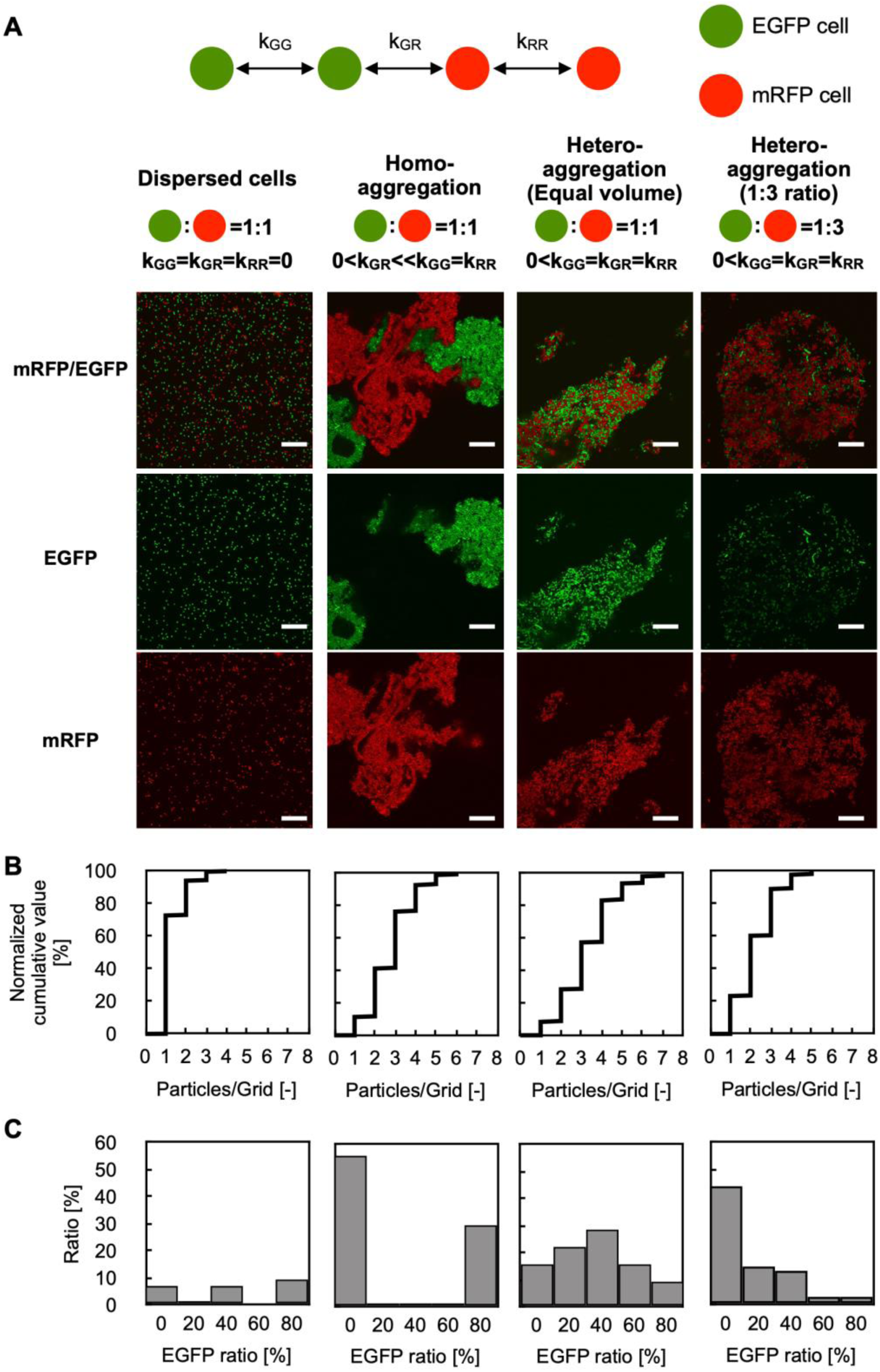
GPIA of dispersed or aggregated cells. (A) CLSM images and schematic cartoons of four control samples: dispersed cells composed of EGFP-AtaA(–) + mRFP-AtaA(–); homo-aggregation obtained by first forming separate EGFP-AtaA(+) and mRFP-AtaA(+) clumps and then mixing them; and hetero-aggregation generated by mixing equal (1:1) or unequal (1:3) volumes of EGFP-AtaA(+) and mRFP-AtaA(+) cells. Scale bars: 20 µm. (B) Frequency distribution of total cell counts in each 2 µm square grid. (C) Ratio of the EGFP-fluorescent cells in each grid containing multiple cells (≥ 1000 cells analyzed per sample).

Next, the ratio of EGFP-fluorescent cells in each grid was calculated (Fig. 3C). In this calculation, particles were integrated as over 1000 events from several image samples because the deviation was almost saturated at over 300 events (Fig. S2). In the dispersed control, no peak was observed because most particles were classified as a single cell in Figure 3B. On the other hand, the homo-aggregation control showed a U-shaped histogram, with high frequencies in the 0–20 % and 80–100 %, indicating that multiple cells of one type are included in each grid. In contrast, the hetero-aggregation control with a 1:1 ratio showed a symmetric unimodal histogram with the peak at 40– 60 %, indicating that two types of multiple cells equally contained in each grid. The hetero-aggregation control with a 1:3 ratio showed a right-skewed histogram, indicating that grids containing one cell type and grids containing two cell types were both present. All pairwise comparisons among the four samples (dispersed cells, homo-aggregates, hetero-aggregates with equal volume, and hetero-aggregates at a 1:3 ratio) showed statistically significant differences in the distribution of the EGFP-ratio, as determined by Pearson’s chi-square test with Bonferroni correction (Table S1). Figure S3 presents the theoretically calculated EGFP-ratio histogram, which displays a distribution closely matching the trend observed in the GPIA-generated histogram. These results demonstrate that GPIA not only distinguishes dispersed from aggregated cell populations but also clearly resolves whether the aggregates arise from homotypic or heterotypic interactions.

### 3.3 Analysis of cell−cell interaction mediated by cells expressing different levels of AtaA

Next, we examined whether the GPIA could detect the difference in the interaction strength between bacterial cells. The mRFP-AtaA(+) cells were cultured in the medium containing 0–0.5 % arabinose to prepare AtaA-displayed cells with different expression levels (Fig. 4A). Immunoblotting showed that the expression level of *ataA* increased according to the arabinose concentration of over 0.05 % (Fig. 4B). Consistent with this, tube-settling assays showed that cells in which AtaA was detectable exhibited self-aggregation, and the aggregation rate increased with higher AtaA levels (Fig. 4C).

**Figure 4.**
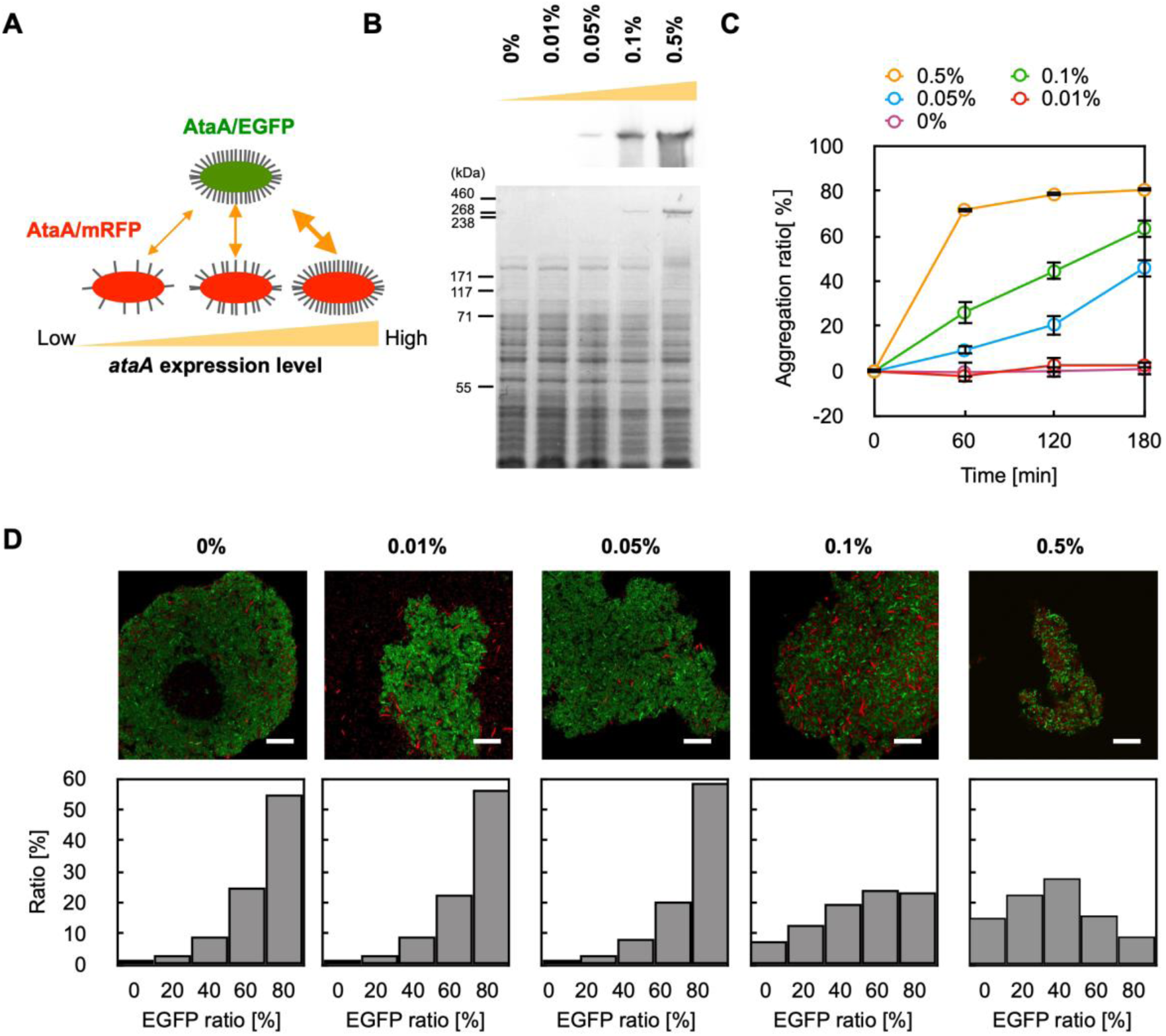
GPIA of co-aggregation mediated by AtaA with different levels of expression. (A) Experimental design: EGFP-AtaA(+) cells induced with 0.5 % arabinose were mixed with mRFP-AtaA(+) cells induced with 0, 0.01, 0.05, 0.1, or 0.5 % arabinose. (B) Immunoblotting and CBB staining of stepwise increases in AtaA production with higher arabinose concentrations. (C) Tube-settling assay of cells induced with 0, 0.01, 0.05, 0.1, or 0.5 % arabinose. The data are presented as the means ± SDs (n = 3). (D) Representative merged CLSM images and corresponding EGFP-ratio histograms calculated by GPIA (≥ 1000 cells per sample). Scale bars: 20 µm.

We then mixed EGFP-AtaA(+) cells grown in 0.5 % arabinose with each mRFP-AtaA(+) grown in 0–0.5 % arabinose and prepared cell aggregates. GPIA revealed that the proportion of hetero-aggregated grids decreased progressively as the AtaA content of the mRFP cells declined (Fig. 4D, Table S2). Notably, cells induced with only 0.05 % arabinose, which expressed little AtaA and aggregated slowly, were scarcely incorporated into the rapidly forming EGFP-AtaA cell aggregates. These results demonstrate that GPIA can detect expression-dependent differences in cell–cell interaction affinity.

### 3.4 Analysis of cell−cell interaction mediated by in-frame deletion mutants of AtaA

Previously, we showed that removing the N-terminal head domain (Nhead) significantly decreased cell adhesion to material surfaces, whereas deleting other parts did not. However, the domain responsible for self-aggregation was still unknown. To identify it, we built cells expressing EGFP and in-frame deletion mutants of AtaA that lack each domain (Fig. 5A) and mixed them with cells expressing mRFP and full-length AtaA (FL-AtaA). We then observed aggregates with CLSM and analyzed them by GPIA (Fig. 5B, C). The cells expressing ΔNS-A1, ΔNS-A2, ΔNS-B, ΔNS-CΔChead, and ΔCstalk formed mixed clumps with cells expressing FL-AtaA and the EGFP-ratio histograms showed a peak at 40–60 %, indicating that aggregates comprised the two types of cells in a 1:1 ratio. In contrast, the ΔNhead mutant failed to co-aggregate—only the cells expressing FL-AtaA aggregated, while the ΔNhead cells stayed dispersed. The EGFP-ratio histogram of ΔNhead showed a peak at 0–20 %, indicating that aggregates were made almost exclusively of mRFP-AtaA(+) cells. A chi-square test showed a significant difference between ΔNhead and the others in the distribution of EGFP-ratio (Table S3). These results indicate that cell–cell interaction mediated by AtaA is driven mainly by homophilic interactions between Nhead of two different cells.

**Figure 5.**
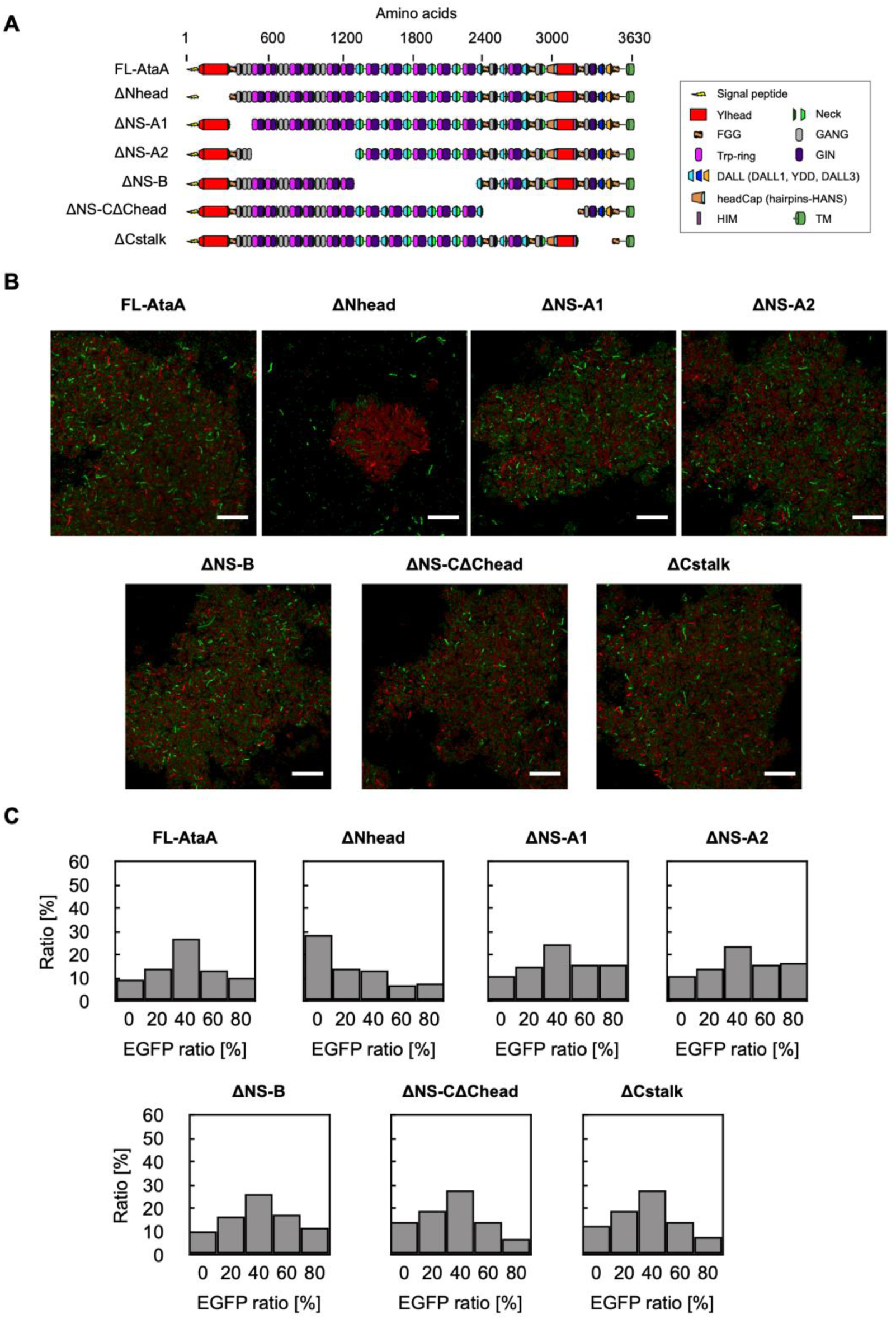
GPIA of cell−cell interaction mediated by in-frame deletion mutants of AtaA. (A) Schematic illustration of full-length AtaA (FL-AtaA) and the in-frame deletion mutants. (B) CLSM images of mixtures containing cells expressing EGFP and AtaA-mutants and cells expressing mRFP and FL-AtaA. Scale bars: 20 µm. (C) EGFP-ratio histograms calculated by GPIA (≥ 1000 cells per sample). A chi-square test confirmed a significant difference between the ΔNhead and the other mutants (see Table S3).

## 4. Discussion

In this study, we developed grid partitioning image analysis (GPIA) that transforms confocal micrographs into quantitative data and resolves both the presence of bacterial aggregates and the composition of the aggregates. The method correctly separated four reference conditions: fully dispersed suspensions, homo aggregates, and two hetero aggregate mixtures (Fig. 3). Because GPIA relies only on standard ImageJ functions and a spreadsheet template, the workflow can be completed in minutes without custom code, making it an easy and accessible method. Compared with classical tube-settling assays that monitor turbidity, GPIA overcomes a key limitation: it can measure interactions among strains that are self-aggregative. By changing AtaA expression with arabinose, we showed that GPIA detects expression-dependent shifts in cell–cell affinity (Fig. 4). The analysis of cells expressing in-frame mutants revealed that Nhead-Nhead interaction is important for the cell aggregation of Tol 5 through AtaA (Fig. 5). These results suggest that GPIA is useful for analyzing adhesin mutants, environmental cues, or inhibitory compounds that produce modest phenotypic changes.

By adjusting the grid size, GPIA can be readily applied to aggregation analyses of bacteria with different dimensions. For instance, a 4-µm grid is likely appropriate for cells with a diameter of about 2 µm (e.g., some species of *Deinococcus*, *Sarcina*, and *Aquisphaera*) (Bondoso et al., 2011; Floc’h et al., 2019; Marcelino et al., 2021). When examining mixed populations of markedly different sizes, such as bacteria and yeast, it may be necessary to re-optimize not only the grid size but also the classification thresholds. In GPIA, a small proportion of physically proximate but non-interacting cells may sometimes be counted as aggregates. This can be minimized by diluting the cell suspension before CLSM observation and analyzing a sufficiently large number of fields. When increasing sample size, it is better to increase the number of observed fields rather than increase cell density because high-concentration samples cause misclassification simply due to spatial crowding. Further statistical analysis of the histogram of EGFP-ratio between samples using Pearson’s chi-square test would help to consider the significant differences.

In conclusion, we developed the GPIA, a rapid, simple, and sensitive method for quantifying bacterial cell aggregation. By converting confocal images into robust numerical data, GPIA bridges the gap between qualitative microscopy and quantitative, yet technically demanding, single-cell analysis. GPIA will accelerate research on cell– cell interactions, which are the basis of important bacterial functions such as surface colonization, tolerance to environmental stress, and interspecies metabolite exchange.

## Data availability

The original contributions presented in the study are included in the article and the supplementary material, further inquiries can be directed to the corresponding author.

## Author contributions

YO: Conceptualization, Formal analysis, Investigation, Methodology, Software, Writing – original draft; SY: Investigation, Writing – original draft, Writing – review & editing; KH: Supervision, Writing – review & editing.

## Acknowledgments

The authors thank Tomoya Karakama for his technical assistance. This work was supported by the Japan Society for the Promotion of Science (JSPS) KAKENHI (Grant Numbers JP21H05227 and JP24H00043).

