## Supplementary Material for "Grid partitioning image analysis for bacterial cell aggregates"

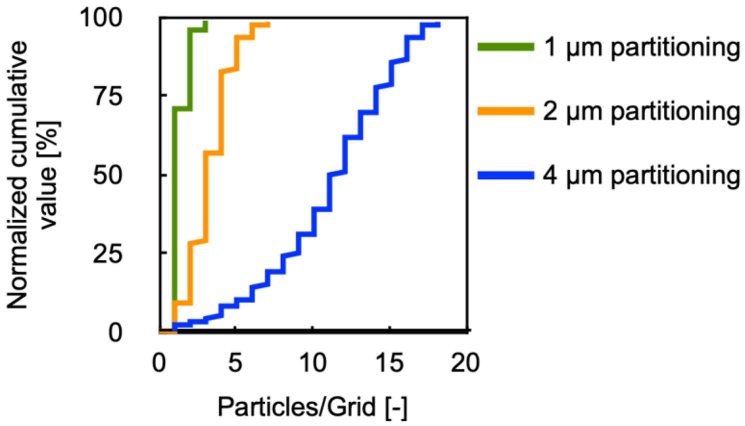


**Figure S1. Particle numbers in each grid.**

The particle counts in each grid was integrated when the partitioning size was set to 1, 2, and 4 µm of a square. This analysis was performed on the hetero-aggregation control images shown in Figure 3A.

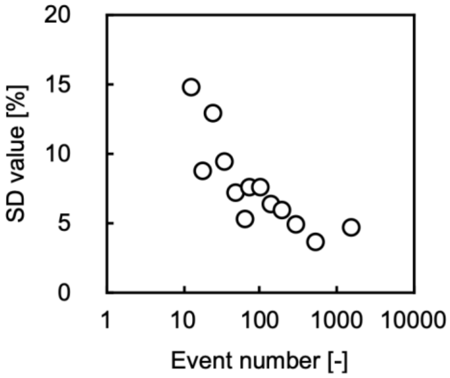


**Figure S2. Calculation of the SD values of EGFP-ratio.**

The horizon axis means the total number of analyzed particles. The vertical axis means the average SD values of each EGFP-ratio in histograms. This analysis was performed on the hetero-aggregation control images shown in Figure 3A.

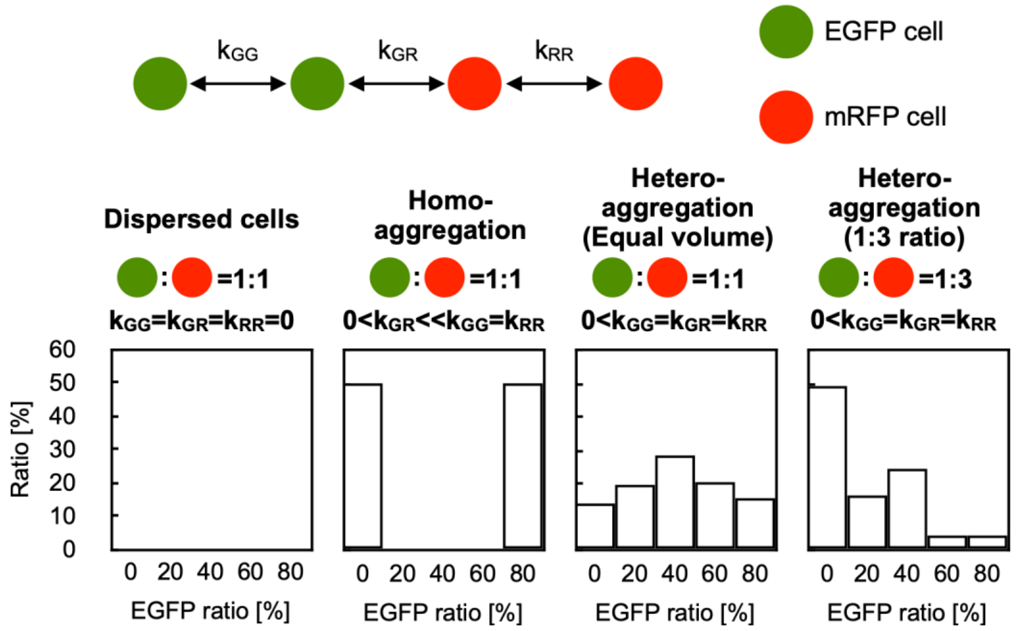


**Figure S3. Theoretical values of EGFP-ratio.**

The theoretical values of EGFP-ratio of each control sample in Figure 3 were calculated.

**Table S1. Statistical comparisons of the EGFP-ratio in Figure 3C by Pearson's chi-square test with Bonferroni correction.**

| | $Chi^2$ | p-value | Cramér's V | 95% CI Lower | 95% CI Upper | Bonferroni-adjusted p |
|---|---|---|---|---|---|---|
| Dispersed vs Homo | 1884 | $< 1.0e^{-300}$ | 0.697 | 0.675 | 0.72 | $< 1.0e^{-300}$ |
| Dispersed vs Hetero (1:1) | 1606 | $< 1.0e^{-300}$ | 0.597 | 0.574 | 0.621 | $< 1.0e^{-300}$ |
| Dispersed vs Hetero (1:3) | 1230 | $8.50e^{-264}$ | 0.518 | 0.493 | 0.543 | $5.10e^{-263}$ |
| Homo vs Hetero (1:1) | 2522 | $< 1.0e^{-300}$ | 0.659 | 0.640 | 0.679 | $< 1.0e^{-300}$ |
| Homo vs Hetero (1:3) | 1816 | $< 1.0e^{-300}$ | 0.555 | 0.534 | 0.577 | $< 1.0e^{-300}$ |
| Hetero (1:1) vs Hetero (1:3) | 1121 | $2.79e^{-240}$ | 0.415 | 0.392 | 0.437 | $1.67e^{-239}$ |

**Table S2. Statistical comparisons of the samples in Figure 4D by Pearson's chi-square test with Bonferroni correction.**

| | Chi$^2$ | p-value | Cramér's V | 95% CI Lower | 95% CI Upper | Bonferroni-adjusted p |
|---|---|---|---|---|---|---|
| 0.5-0.5% vs 0.5-0.1% | 1045 | $7.01e^{-224}$ | 0.175 | 0.165 | 0.186 | $7.01e^{-223}$ |
| 0.5-0.5% vs 0.5-0.05% | 5891 | $< 1.0e^{-300}$ | 0.448 | 0.438 | 0.458 | $< 1.0e^{-300}$ |
| 0.5-0.5% vs 0.5-0.01% | 5086 | $< 1.0e^{-300}$ | 0.495 | 0.483 | 0.507 | $< 1.0e^{-300}$ |
| 0.5-0.5% vs 0.5-0% | 7015 | $< 1.0e^{-300}$ | 0.375 | 0.367 | 0.383 | $< 1.0e^{-300}$ |
| 0.5-0.1% vs 0.5-0.05% | 8069 | $< 1.0e^{-300}$ | 0.376 | 0.368 | 0.383 | $< 1.0e^{-300}$ |
| 0.5-0.1% vs 0.5-0.01% | 5774 | $< 1.0e^{-300}$ | 0.345 | 0.336 | 0.353 | $< 1.0e^{-300}$ |
| 0.5-0.1% vs 0.5-0% | 9756 | $< 1.0e^{-300}$ | 0.354 | 0.348 | 0.361 | $< 1.0e^{-300}$ |
| 0.5-0.05% vs 0.5-0.01% | 50 | $1.08e^{-09}$ | 0.032 | 0.023 | 0.041 | $1.08e^{-08}$ |
| 0.5-0.05% vs 0.5-0% | 188 | $6.69e^{-39}$ | 0.050 | 0.043 | 0.057 | $6.69e^{-38}$ |
| 0.5-0.01% vs 0.5-0% | 39 | $2.01e^{-07}$ | 0.023 | 0.015 | 0.031 | $2.01e^{-06}$ |

**Table S3. Statistical comparisons of the samples in Figure 5C by Pearson's chi-square test with Bonferroni correction.**

| | Chi$^2$ | p-value | Cramér's V | 95% CI Lower | 95% CI Upper | Bonferroni-adjusted p |
|---|---|---|---|---|---|---|
| ΔNhead vs FL-AtaA | 2976 | $< 1.0e^{-300}$ | 0.251 | 0.242 | 0.259 | $< 1.0e^{-300}$ |
| ΔNhead vs ΔNS-A1 | 2722 | $< 1.0e^{-300}$ | 0.287 | 0.277 | 0.297 | $< 1.0e^{-300}$ |
| ΔNhead vs ΔNS-A2 | 2857 | $< 1.0e^{-300}$ | 0.282 | 0.272 | 0.292 | $< 1.0e^{-300}$ |
| ΔNhead vs ΔNS-B | 2722 | $< 1.0e^{-300}$ | 0.314 | 0.302 | 0.325 | $< 1.0e^{-300}$ |
| ΔNhead vs ΔNS-CΔChead | 1974 | $< 1.0e^{-300}$ | 0.252 | 0.241 | 0.263 | $< 1.0e^{-300}$ |
| ΔNhead vs ΔCstalk | 2423 | $< 1.0e^{-300}$ | 0.262 | 0.252 | 0.272 | $< 1.0e^{-300}$ |

**File S1. Template spreadsheet for GPIA.**

Input the coordinates of EGFP and mRFP. The histogram of EGFP-ratio is output on the right side of the spreadsheet. A spreadsheet containing model data is also included.
